# Development and validation of an expanded carrier screen that optimizes sensitivity via full-exon sequencing and panel-wide copy-number-variant identification

**DOI:** 10.1101/178350

**Authors:** Gregory J. Hogan, Valentina S. Vysotskaia, Stefanie Seisenberger, Peter V. Grauman, Kyle A. Beauchamp, Kevin R. Haas, Sun Hae Hong, David Jennions, Diana Jeon, Shera Kash, Henry H. Lai, Laura M. Melroy, Mark R. Theilmann, Clement S. Chu, Saurav Guha, Kevin Iori, Jared R. Maguire, Kenny K. Wong, Eric A. Evans, Imran S. Haque, Rebecca Mar-Heyming, Hyunseok P. Kang, Dale Muzzey

**Affiliations:** Counsyl, Inc., South San Francisco, CA, USA; Current affiliation: Freenome, Inc., South San Francisco, CA, United States

**Keywords:** expanded carrier screening, genetic testing, next-generation sequencing, analytical validation

## Abstract

**Purpose:** By identifying pathogenic variants across hundreds of genes, expanded carrier screening (ECS) enables prospective parents to assess risk of transmitting an autosomal recessive or X-linked condition. Detection of at-risk couples depends on the number of conditions tested, the diseases’ respective prevalences, and the screen’s sensitivity for identifying disease-causing variants. Here we present an analytical validation of a 235-gene sequencing-based ECS with full coverage across coding regions, targeted assessment of pathogenic noncoding variants, panel-wide copy-number-variant (CNV) calling, and customized assays for technically challenging genes.

**Methods:** Next-generation sequencing, a customized bioinformatics pipeline, and expert manual call review were used to identify single-nucleotide variants, short insertions and deletions, and CNVs for all genes except *FMR1* and those whose low disease incidence or high technical complexity precludes novel variant identification or interpretation. Variant calls were compared to reference and orthogonal data.

**Results:** Validation of our ECS data demonstrated >99% analytical sensitivity and >99% specificity. A preliminary assessment of 15,177 patient samples reveals the substantial impact on fetal disease-risk detection attributable to novel CNV calling (13.9% of risk) and technically challenging conditions (15.5% of risk), such as congenital adrenal hyperplasia.

**Conclusion:** Validated, high-fidelity identification of different variant types—especially in diseases with complicated molecular genetics—maximizes at-risk couple detection.

## INTRODUCTION

There are more than 1000 recessive single-gene conditions that vary in both severity and age of onset.^1^ Each is uncommon in the general population, yet collectively these Mendelian diseases account for approximately 20% of infant mortality and 10% of infant hospitalizations.^2,3^ Screening for carriers of such conditions in the preconception or prenatal period informs couples about both the risk of having a child with a serious disease and the available family-planning options. The risk assessment and patient autonomy provided by carrier screening can have substantial impact, as nearly 2 million women give birth to their first child each year in the United States.^4^

Due to the rising quality and falling cost of genomic technologies, it is now possible to perform pan-ethnic carrier screening on a large number of conditions simultaneously (referred to as “expanded carrier screening”, or “ECS”). Our recent retrospective study of carrier rates in 346,790 patients showed that an ECS panel is expected to identify more pregnancies at risk for severe or profound conditions than ethnic-based panels spanning far fewer genes.^5^ Citing this work, the American College of Obstetricians and Gynecologists (ACOG) issued new guidelines in March 2017, recognizing expanded carrier screening as an acceptable strategy for prepregnancy and prenatal carrier screening for patients and their partners.^6^

An ECS must have a high detection rate for each disease on the panel to identify at-risk couples and to minimize the residual risk in couples where only one partner has tested positive. Indeed, detection rate is particularly important for the recessive diseases that predominate ECS panels because the odds of detecting an at-risk couple scales as the square of the rate for finding an individual carrier (e.g., 80% detection rate for one parent means only a 64% detection rate for an at-risk couple). Relative to the targeted genotyping approaches used for classical carrier screening, next-generation sequencing (NGS) has enabled ECS panels to achieve very high per-disease detection rates, as both common and rare variants can be identified across the entire coding region and in relevant noncoding positions of the disease gene.^1,7–11^ Underscoring the at-risk couple detection gain afforded by NGS, a study of 11,691 individuals screened for 15 genes by NGS revealed that approximately one quarter carried mutations not typically included in targeted panels.^8^

To maximize detection rates, novel copy number variants (CNVs) must be identified, yet most ECS offerings—even those using NGS—report only single nucleotide variants (SNVs), indels, and, at most, a small handful of common CNVs with known breakpoints. However, CNVs can vary in size and position, encompassing everything from single exons (which account for 29% of CNVs for Mendelian conditions^12^) to the entire gene. A diversity of pathogenic CNVs has been observed in cystic fibrosis carriers, accounting for 1.6% of carriers^13^, meaning that the single-carrier detection rate without CNV detection is <98.4%, which in turn makes the at-risk couple rate <96.8%. The inverse is noteworthy: including novel CNV detection can boost at-risk couple detection for cystic fibrosis to nearly 100%. To our knowledge, the impact of CNVs across all genes on an ECS has not yet been characterized.

Carrier status for a minority of the most prevalent serious conditions is difficult to resolve with standard NGS and bioinformatics approaches due to the challenging sequence features of the disease gene; thus, these conditions require special handling. Low complexity spans (e.g., CGG repeat expansion in *FMR1* for fragile X syndrome) and highly homologous regions (e.g., *SMN1* and *SMN2* genes for spinal muscular atrophy) complicate variant identification, yet these hard-to-sequence genes simultaneously contribute substantially to the disease risk. For instance, fragile X syndrome, spinal muscular atrophy, 21-hydroxylase deficient congenital adrenal hyperplasia (21-OH CAH), and alpha thalassemia account for 54 affected fetuses per 100,000 pregnancies.^14^

Here, we validate and describe an ECS (Counsyl Foresight™ Carrier Screen) leveraging NGS to identify SNVs, indels, novel CNVs (deletions for nearly all genes and both deletions and duplications for *CFTR* and *DMD*), and hard-to-sequence targeted variants. Following recommendations of the College of American Pathologists (CAP)^15^ and the American College of Medical Genetics and Genomics (ACMG)^16^, we measured the analytical accuracy, precision, sensitivity, and specificity of the test. Further, on a cohort of 15,177 patients, we performed a preliminary assessment of the impact of panel-wide CNV calling on detection of at-risk pregnancies. Our data demonstrate the accurate detection of the most common types of genomic alterations found in reference cell lines and routine clinical specimens.

## MATERIALS AND METHODS

### Institutional Review Board approval

The protocol for this study was approved by Western Institutional Review Board (IRB number 1145639) and complied in accordance with the Health Insurance Portability and Accountability Act (HIPAA). The information associated with patient samples was de-identified in accordance with the HIPAA Privacy Rule. A waiver of informed consent was requested and approved by the IRB.

### Test description

We compiled a panel (Counsyl Foresight™ Carrier Screen) of 235 genes responsible for 234 clinically important autosomal recessive and X-linked diseases (**Table S1**). This panel consists of a “Universal” sub-panel (176 diseases) for routine ECS and an opt-in panel (234 diseases) aimed at specific high-risk populations. The design of the Universal panel is described in Beauchamp et al^14^ and prioritized diseases that are prevalent, with serious and highly penetrant phenotypes that would likely affect clinical counseling for preventative measures and family planning. Some genes for high-prevalence diseases with moderate but lifelong impact are also included (e.g., *MEFV* and *GJB2*). The gene-specific methodologies and types of variants reported are summarized in **Table 1**.

**Table 1.** Foresight™ expanded carrier screening panel

| GENERAL ECS |  |  |
| --- | --- | --- |
| Disease Genes | Methodology | Variants Reported |
| 216 genes | NGS | Novel pathogenic SNVs, indels, large deletions |
| *CFTR* | NGS | Novel pathogenic SNVs, indels, large deletions and duplications |
| *DMD* | NGS | Novel pathogenic SNVs, indels, large deletions and duplications |
| 11 genes | NGS | Targeted pathogenic mutations (**Table S1**) |
| TECHNICALLY CHALLENGING GENES |  |  |
| Disease Genes | Methodology | Variants Reported |
| *SMN1* | NGS | Exon 7 copy number, g.27134T>G SNP |
| *CYP21A2* | NGS | **Classical:** *CYP21A2* 30kb deletion, *CYP21A2* duplication, *CYP21A2* triplication, c.293-13C>G, p.G111Vfs*21, p.I173N, p.[I237N;V238E;M240K], p.L308Ffs*6, p.Q319*, p.Q319* + *CYP21A2*dup, p.R357W<br>**Non-classical:** p.P31L, p.V281L |
| *HBA1/2* | NGS | **Single deletions:** -alpha3.7, -alpha4.2<br>**Double deletions:** -(alpha)20.5, --BRIT, --MEDI, --MEDII, --SEA, --THAI or --FIL<br>**Frequent SNV:** Hb Constant Spring<br>**Regulatory deletion:** ΔHS-40<br>(combinations of most variants above with other deleterious variants or duplications can also be detected) |
| *GBA* | NGS | p.N370S, p.D409V, p.D448H, IVS2+1G>A, p.L444P, p.R463C, p.R463H, p.R496H, p.V394L, p.L29Afs*18 |
| *FMR1* | PCR/CE | Number of CGG repeats in the 5' UTR |

PCR/CE, polymerase chain reaction/capillary electrophoresis; UTR - untranslated region

### Next generation sequencing, bioinformatics processing, and variant interpretation

#### Next generation sequencing

The molecular workflow of our NGS pipeline was previously described.^17^ Briefly, DNA is fragmented to 200-1000 bp by sonication and then converted to a sequencing library by end repair, A-tailing, and adapter ligation. Samples are then amplified by PCR with barcoded primers, multiplexed, and subjected to hybrid capture-based enrichment with 40-mer oligonucleotides. Sequencing of the selected targets is performed on the Illumina HiSeq 2500 instrument.

#### Bioinformatics processing

Sequencing reads are aligned to the hg19 human reference genome using the BWA-MEM algorithm.^18^ Novel SNVs and indels are identified and genotyped using GATK 1.6 and FreeBayes^19,20^, and nine known-pathogenic sites involving complex indels are detected with custom genotyping software. Copy number variants are determined using custom software that leverages read-depth values.^17^ A combination of targeted genotyping and read depth-based copy number analysis is used to determine the number of functional gene copies and/or the presence of selected mutations in technically challenging genes, such as *SMN1*, *GBA*, *HBA1/2*, and *CYP21A2* (**Supplementary Methods**).

#### Quality control metrics

Ancillary quality-control (QC) metrics are computed on the sequencing output and used to exclude and re-run failed samples. QC metrics include the fraction of sample contamination (<5%), extent of GC bias, read quality (percent Q30 bases per Illumina specifications), depth of coverage (mean coverage of >50x), and region of interest (ROI) coverage (>99% per base minimum coverage >=20x) (**Table S2**). Calls that do not meet QC criteria are set to “no-call”.

#### Variant review and interpretation

To ensure clinical calling accuracy, all calls and no-calls for pathogenic, likely pathogenic, and uncurated variants are manually reviewed by laboratory personnel and are subject to override if warranted, based on a pre-established protocol. Identified variants are classified according to the ACMG Standards and Guidelines for the Interpretation of Sequence Variants^21^ as described previously.^17^ Final variant classifications are regularly uploaded to ClinVar (National Center for Biotechnology Information; https://www.ncbi.nlm.nih.gov/clinvar/).^22^

### *FMR1* CGG repeat sizing

CGG trinucleotide expansions of the *FMR1* promoter are measured by PCR amplification and capillary electrophoresis as previously described.^23^

### Analytical validation

*Samples and reference data.* Samples and reference data are compiled from different sources (**Tables S3 and S4**). Purified DNA for 91 cell lines from the 1000 Genomes (1KG) Project^24^ and 70 cell lines with known pathogenic variants in specific genes were purchased from the Coriell cell line repository (Camden, NJ) (**Tables S5A-E**). In addition, 115 mutation-positive patient blood and saliva samples tested with a previous version of the Counsyl carrier test (a 94-gene panel) were included in the validation. Relevant variants in all mutation-positive patient samples were confirmed orthogonally by PCR/Sanger sequencing, quantitative PCR, or multiplex ligation-dependent probe amplification (MLPA). Further details on confirmatory testing can be found in the **Supplementary Methods**. Note that NA06896 was dropped from *FMR1* accuracy analysis due to inconsistent reference data.^25,26^

*1KG analysis.* Ninety-one 1KG samples were sequenced to measure the accuracy of SNV and short indel calls in 229 genes (**Table S5A**). Ninety samples passed QC and manual review. *Simulation of synthetic CNVs.* For every region reportable for CNVs, we simulated a single-copy deletion and tested calling sensitivity; we also simulated single-copy duplications for *DMD* and *CFTR* regions (detailed description of CNV simulations in the **Supplementary Methods**). *Statistical analysis.* Validation metrics were defined as: Accuracy = (TP + TN) / (TP + FP + TN + FN); Sensitivity = TP / (TP + FN); Specificity = TN / (TN + FP); FDR = FP / (TP + FP), where TP - true positives, TN - true negatives, FP - false positives, FN - false negatives, and FDR - false discovery rate. The confidence intervals (CIs) were calculated by the method of Clopper and Pearson.^27^ Reproducibility within and between runs was calculated as the ratio of concordant calls to total calls.

## RESULTS

We developed an NGS-based ECS (Counsyl Foresight™ Carrier Screen) covering 220 autosomal recessive and 14 X-linked conditions, including technically challenging diseases (**Figure 1**, **Tables 1 and S1**). For nearly all genes on the Universal panel, SNVs and indels are detected via NGS data acquired from regions that could impact gene function (e.g., padded coding exons and known or potentially pathogenic intronic variants; **Figure 1**, **top left**). Large CNVs are identified at single-exon resolution panel-wide using relative sample-to-sample changes in sequencing depth (**Figure 1**, **top right;** see **Methods**). The test has been validated (described in detail below) and used in a clinical production setting on 15,177 patient samples (tested between Nov. 2016 and Aug. 2017). Carrier rates on this cohort enable calculation of the expected fraction of US pregnancies whose affected status would be identified by the Universal panel of the ECS on a per-disease level **(Figure 1 middle**): the estimate is that 1 in 300 pregnancies would be affected by at least one serious disease on our ECS. For a handful of prevalent diseases that comprise 8.8% of the total panel disease risk, high carrier sensitivity requires customized CNV analysis due to the genes’ complicated technical features (**Figure 1**, **bottom**).

**Figure 1.**
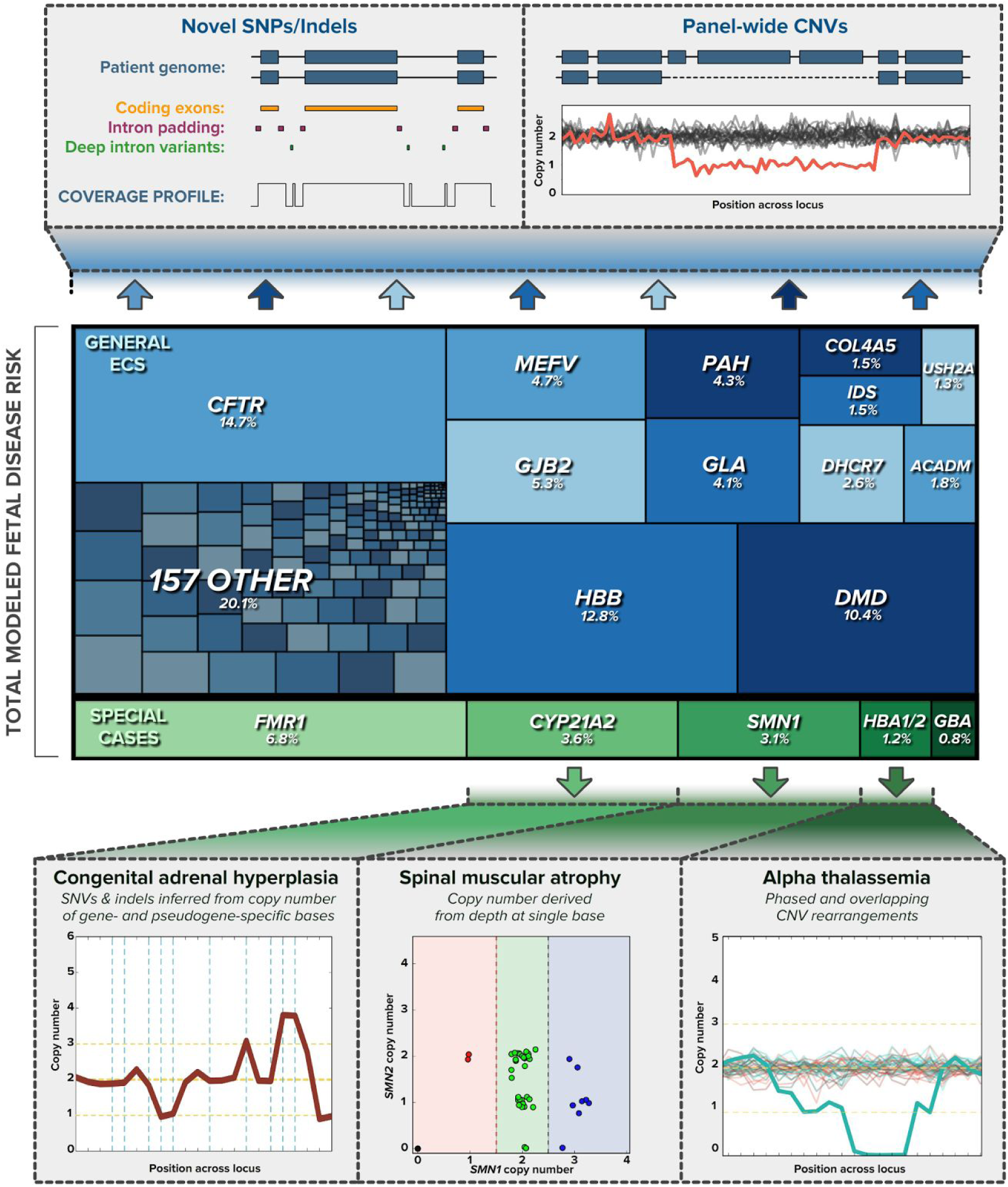
Elements of 176-gene expanded carrier screen that boost detection of at-risk couples. (top) Coverage across padded coding regions—plus known pathogenic intronic sites—together with panel-wide copy-number variant (CNV) calling (positive sample shown in red on background of negative samples in black) identifies couples at risk on the Universal panel (see **Methods**). **(middle)** The modeled fetal disease risk is shown for each disease gene, estimated from carrier rates of 15,177 patients (percent indicates share of total panel disease risk). **(bottom)** Five conditions require special-case treatment, with four leveraging customized CNV calling (see **Supplementary Methods**). For 21-OH deficient congenital adrenal hyperplasia (bottom left) and alpha thalassemia (bottom right), copy-number profiles are plotted from 5’ to 3’ across the gene; for spinal muscular atrophy (bottom middle), each spot represents the copy number of *SMN1* and *SMN2* for a single sample.

Relative to our prior version of the ECS characterized in a recent study of 346,790 patients^5,14^, this updated ECS differs in two ways that collectively boost the assessed fetal disease risk. The updated Universal panel ECS probes SNVs and indels for 82 more diseases (**Figure 2A,B**), and it additionally detects deletions ranging in size from a single exon to the entire gene (**Figure 2B,C**). Panel-wide CNVs contribute approximately 14% of the assessed fetal disease risk: CNVs in *DMD* alone represent ∼10% of the risk, with the remaining genes accounting for ∼4%, largely consistent with our estimate of CNV-attributable disease risk estimated from the previous 94 disease panel.^14^

**Figure 2.**
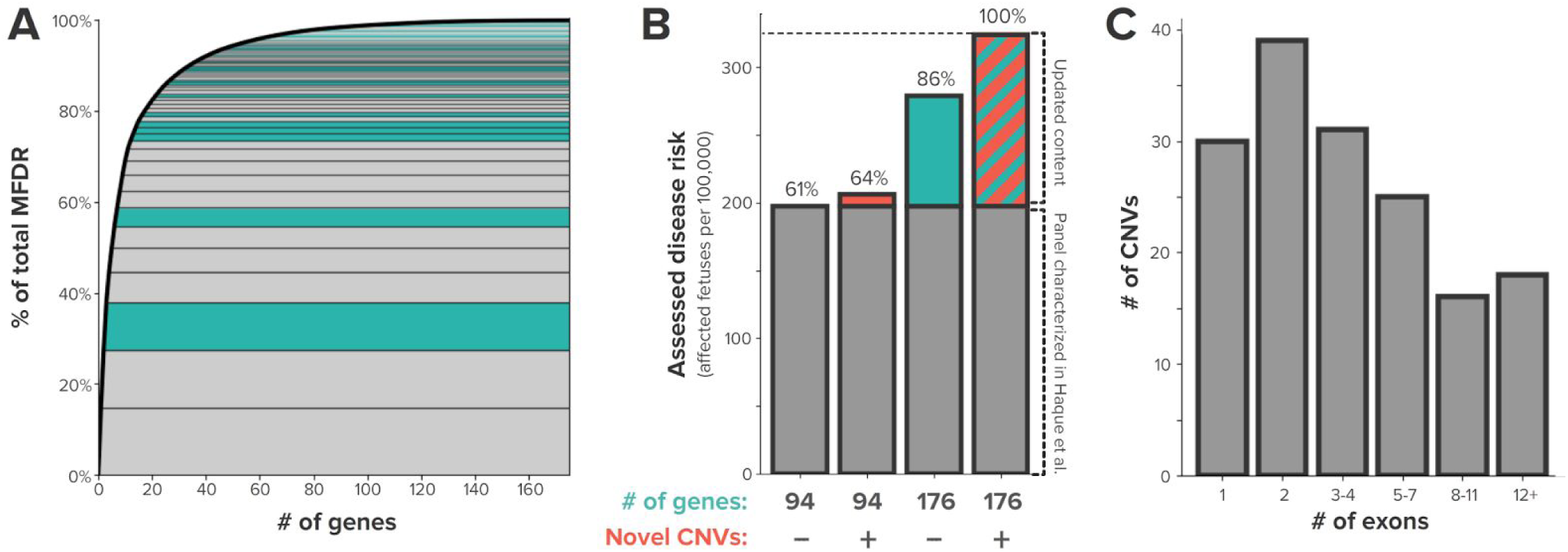
Gain in detected affected fetuses resulting from panel expansion and novel copy-number variant calling. **(A)** For the 176 severe or profound conditions on the Universal panel of our expanded carrier screen applied to 15,177 clinical samples, the relative contribution to the modeled fetal disease risk (MFDR) of each gene (green for genes added in panel update) is plotted as a cumulative distribution. **(B)** Relative contributions to assessed disease risk of 82 additional disease genes and novel CNV calling in the Universal panel. **(C)** The size distribution, expressed in exons, of observed deletions.

### Validation approach

To assess the analytical performance of the ECS panel prior to launching it in a clinical setting, we measured the accuracy of identifying variants that are small (e.g., SNVs and small indels), technically nuanced (e.g., large indels and CNVs), and in hard-to-sequence genes (e.g., *CYP21A2*) (**Table S4)**. Our validation approach builds upon the broad recommendations from the College of American Pathologists (CAP)^15^ and the American College of Medical Genetics and Genomics (ACMG)^16^ for validation of targeted, germline testing using NGS.

Validation of a new assay requires a reference set for comparison. For smaller-sized variant types, reference data and samples are readily available; however, for technically challenging genes and variants, reference material is scarce. To gather reference data that tests such challenging variants, we identified relevant patient samples tested with a previous version of the Counsyl ECS (a 94-gene panel) and orthogonally confirmed each positive variant by a Sanger, TaqMan, or MLPA assay, as appropriate. Since we also wanted to establish CNV-calling proficiency in regions for which no reference samples exist, we implemented *in silico* simulations of CNVs.

Collectively, the validation reference dataset establishes a high standard for validating ECS panels by spanning both the scope of clinical sample types (**Table S3**) and the range of variant types (**Table S4**).

### Accuracy and reproducibility for calling SNPs and small indels in 229 genes

Validation samples were tested using our standard operating procedure (SOP), which includes in- and post-process QC at the batch, sample, and variant-call level (**Table S2**). Furthermore, consistent with our SOP for clinical samples, licensed experts, who were blind to the validation sample set, performed manual review of the sequencing data using our custom review interface. Samples that failed QC and manual review were excluded from further analysis.

We compared our ECS data for the reference sample NA12878 with data from the Genome in a Bottle Consortium^28^, which includes high-confidence calls for >97.5% of the regions covered by our test. We tested NA12878 across five batches and in duplicate within three batches for a total of eight tests, and the test results were highly accurate (>99.99%, **Table S6**). As NA12878 is one of our routine controls within every production batch, since the validation of our test we have further measured accuracy across 207 batches that span reagent lots and instruments, consistently observing high calling accuracy across the panel (**Figure S1**).

To measure SNV and indel calling accuracy across a diverse set of samples, we performed our ECS on 1KG samples. For 90 samples that we tested and passed QC, we compared genotypes across all exonic regions with sufficient coverage and quality in the 1KG data (248,490 calls in all); 52 discordant calls were adjudicated with Sanger sequencing (**Table S7**). Our ECS identified 36,032 true-positive calls and 212,139 true-negative calls, resulting in >99.99% accuracy, sensitivity, and specificity (**Figure 3A**).

**Figure 3.**
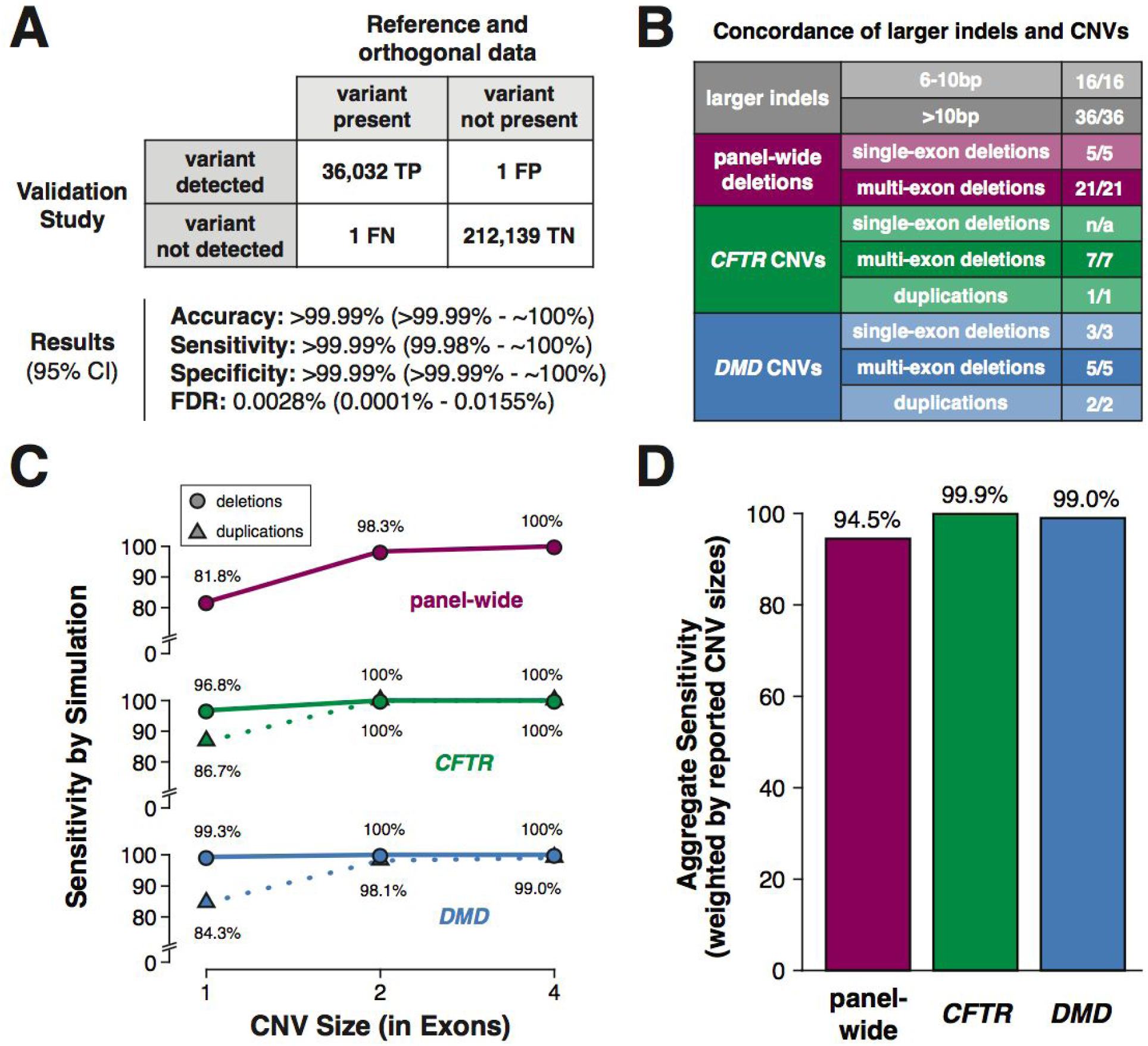
Analytical performance for calling SNVs, indels, and CNVs. **(A)** Contingency table and results for SNV and small indel calling in 229 genes, assessed using 1KG reference material and adjudication by follow-up Sanger. For true-negative calculations, all polymorphic positions (positions at which we observed non-reference bases in any sample) across all samples were considered. No-calls were censored from analysis. The no-call rate was 0.13% (317 / 248,490). CI - confidence interval, TP - true positives, TN - true negatives, FP - false positives, FN - false negatives, and FDR - false discovery rate. **(B)** Concordance summary for larger indels and copy number variants (CNVs). **(C)** Sensitivity for CNV calling as measured by simulations, by gene, type, and size (in number of exons). **(D)** Aggregate sensitivity for CNV calling as measured by simulations. Simulation results in (C) were weighted by size and frequency (see Supplementary Methods). In (B-D), data reported for “panel-wide” deletions exclude *CFTR* and *DMD*.

In addition to establishing the analytical accuracy of the ECS using reference DNA from cell lines, we measured intra- and inter-run reproducibility using different sample types by comparing the equivalence of genotyping calls starting from separate aliquots of DNA. Overall, the test achieved >99.9% intra- and inter-assay reproducibility (**Table S8**).

### Technically challenging variants

#### Larger indel detection performance

Although only 5% of indels are ≥5bp^29^, sensitivity falls as indel size grows. Thus, to ensure high analytical sensitivity for detecting indels, we built a cohort of 52 patient samples with 49 unique technically-challenging, larger (>5bp) deletions, insertions, or complex indels in 42 different genes (**Table S9A**). All of the expected indel calls (52/52), including a 33bp deletion and 21bp insertion, were observed (**Figure 3B**).

#### CNV detection performance

To overcome the limitation of scarce reference materials for CNV calling, we supplemented available reference material with orthogonally confirmed positives identified retrospectively and additionally used *in silico* simulated CNVs to measure sensitivity systematically across the panel. In the empirical analysis, the reference set included 11 Coriell cell lines with a known CNV (**Table S5B**) and 33 clinical samples with CNVs that we confirmed by MLPA (**Table S9B**). The 44 CNV-positive samples included 41 deletion variants in 13 different genes and 3 duplication variants in *CFTR* and *DMD*. Notably, 23 samples had a single-exon or two-exon CNV, which can be technically challenging for a NGS-based assay (**Table S4**). We assessed CNV calling performance separately for *DMD* and *CFTR*, for which we optimized read depth to ensure high sensitivity for both deletions and duplications at the single-exon level. As shown in **Figure 3B**, we detected all 44 CNVs, demonstrating high sensitivity (100%; 95% CI, 92%-100%; reproducibility data in **Table S10**).

In the *in silico* CNV simulations, we introduced a synthetic deletion or duplication spanning at least one coding exon in the background of empirical sample data from four validation flowcells (see **Supplementary Methods**). To assess our ability to call single-exon and multi-exon CNVs, three categories of synthetic CNVs (one-, two-, and four-exon blocks) were tested, with each synthetic CNV size and position being simulated independently using 20 different samples as background (yielding >250k total simulations). Results are summarized in **Figure 3C,D**. To assess sensitivity of clinically relevant deletions and duplications in *CFTR* and *DMD*, we scaled the sensitivity for each CNV size by its population frequency cataloged in public databases (The Clinical and Functional TRanslation of CFTR (CFTR2), https://www.cftr2.org; the Leiden Open Variant Database (LOVD)^30^, http://www.dmd.nl), yielding an aggregate 99.9% sensitivity for CNVs in each gene. Across the rest of the panel, for which only deletions are reported, our simulations revealed 81.8% sensitivity for single-exon deletions and 98.3%-100% sensitivity for multi-exon deletions. Taken together, the simulation results suggest our ECS has high proficiency in identifying CNVs with exon-level resolution.

#### Variant detection performance using NGS in the technically challenging genes *CYP21A2*, *HBA1/2*, *GBA*, and *SMN1*

Several diseases of clinical importance result from mutations in genes that have a paralog or pseudogene that complicates molecular analysis. Such diseases include spinal muscular atrophy (*SMN1* and *SMN2* encode the same protein, but *SMN2* harbors a splicing variant that results in ∼10% of functional SMN protein relative to *SMN1*^*31*^), alpha-thalassemia (*HBA1* and *HBA2* have identical coding sequences and few distinguishing noncoding bases), 21-OH deficient CAH (the *CYP21A2* coding sequence is >99% identical to its pseudogene *CYP21A1P*), and Gaucher disease (*GBA* has a nearby pseudogene *GBAP1* with which it shares high sequence identity in certain exons). Recombination and gene conversion is frequent among these genes and their homologs, which can result in copy number changes. To detect deleterious variants in these technically challenging genes, we implemented custom variant-calling algorithms combining depth-based copy number and specific mutation analyses for the disease genes and their homologs (see **Supplementary Methods**). Below, we describe results for measuring sensitivity; a summary of reproducibility can be found in **Table S10**.

#### *CYP21A2* CNV analysis

Mutations in *CYP21A2* account for 3.6% of the risk assessed by our ECS (**Figure 1**), with approximately 65-70% arising from gene conversion events with a pseudogene and 25-30% from large gene rearrangements.^32^ To assess the accuracy of our NGS-based assay to detect large and small *CYP21A2* rearrangements, we tested 14 specimens previously called positive on the Counsyl 94-gene panel (**Table S9C**). We confirmed the variants using MLPA or long-range PCR and Sanger sequencing (see **Supplementary Methods**). All 14 validation samples—whose variants account for >95% of deleterious *CYP21A2* mutations^32^—were genotyped correctly (see **Figure 4A**).

**Figure 4.**
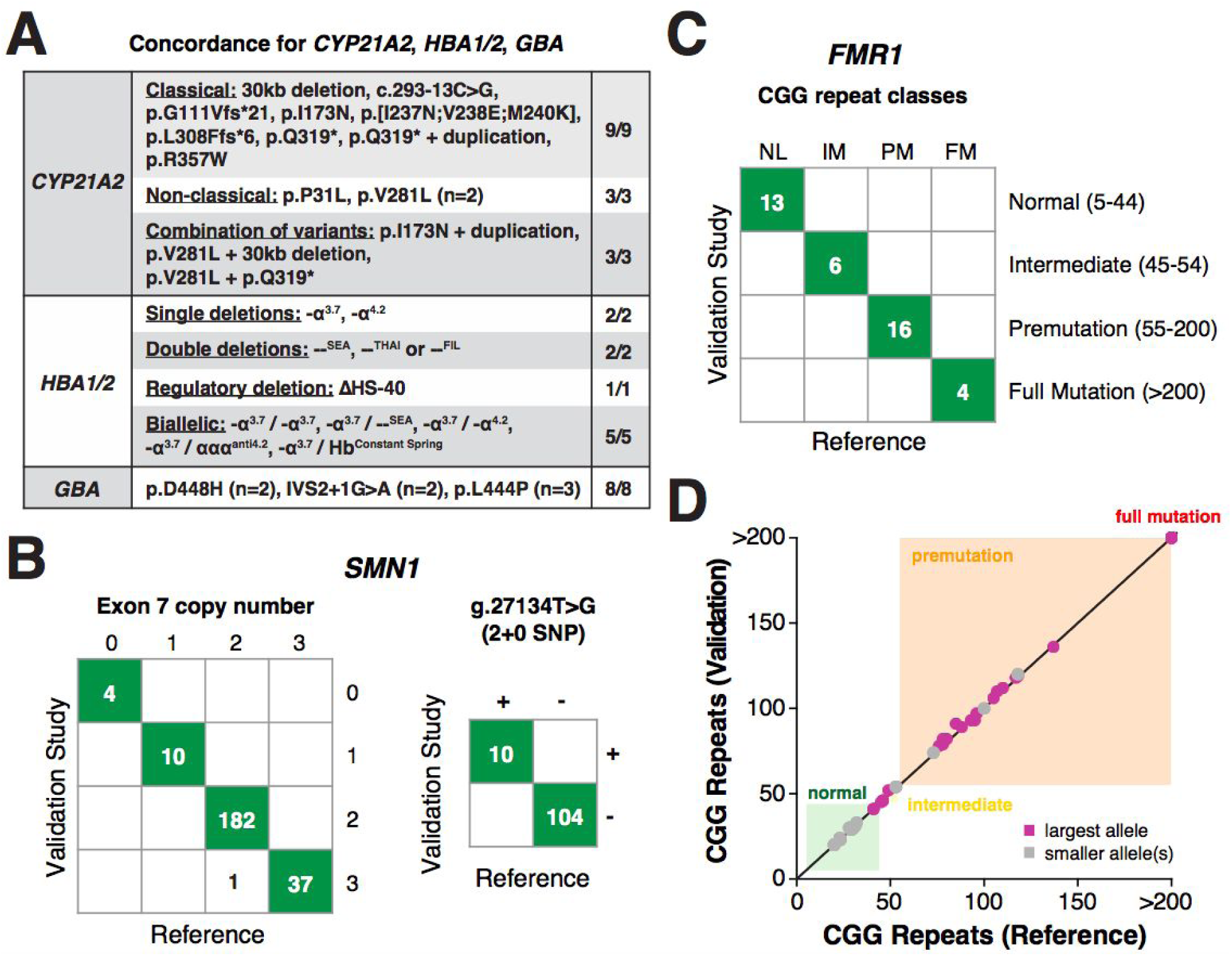
Analytical performance for variant calling in challenging genes. **(A)** Table summarizing concordance for *CYP21A2*, *HBA1/2*, and *GBA*. Unless otherwise stated, one unique sample was tested for each variant listed. **(B)** Concordance of *SMN1* exon 7 copy number calling performance via targeted sequencing and for identification of g.27134T>G, which is associated with silent carriers. **(C)** Concordance for calling FMR1 CGG repeat size, binned into ACMG-defined allele classes. **(D)** Comparison of CGG repeat sizes in the validation study versus the literature consensus for reference cell lines.

#### *HBA1/2* common-variant analysis

Alpha thalassemia represents 1.2% of risk assessed by our ECS (**Figure 1**), and much of that risk arises from deletions among *HBA1* and *HBA2*, which have identical coding sequences. Screening for alpha thalassemia requires detection of the total copy number of *HBA* genes, as well as their phasing on the chromosome. We identified 10 patient samples that had been tested on our previous panel and were found to have mutations in the *HBA1* and/or *HBA2* genes (**Table S9C**). Heterozygous and homozygous deletions for single genes, cis deletions of both genes, and combinations thereof were confirmed during validation (**Figure 4A**).

#### *GBA* CNV analysis

Gaucher disease, which accounts for 0.8% of the total panel disease risk, can arise from gene-conversion events where pseudogenic sequence in *GBAP1* recombines into the *GBA* locus. To validate that our ECS can identify deleterious gene-conversion alleles, six patient samples and one Coriell DNA sample with the pathogenic mutations L444P, D448H, and IVS2+1G>A in *GBA* were successfully identified (**Figure 4A**).

#### *SMN1* copy number and g.27134T>G SNP analyses

Risk for spinal muscular atrophy—3.1% of ECS panel disease risk (**Figure 1**)—is mostly influenced by the copy number of *SMN1*, which is distinguished from the highly homologous *SMN2* gene by an intronic variant that influences splicing of exon 7. To determine the accuracy of NGS-based *SMN1* copy-number calling, 128 unique (234 total with replicates) DNA samples with 0, 1, 2, or 3 copies of *SMN1* were analyzed by NGS (**Tables S4** and **S5D**). Carrier (samples with 0 or 1 copy) versus non-carrier (samples with 2 or more copies) identification accuracy by NGS was 100% (95% CI, 98.4%-100%). NGS copy-number accuracy was 233/234, 99.6% (95% CI, 97.6%-100%) (**Figure 4B**), where one non-carrier patient sample had three copies by NGS and two by TaqMan.

We also measured detection of the g.27134T>G SNP associated with 2+0 SMA carrier status.^33^ The analysis included 98 (92 unique) 1KG cell line samples containing the g.27134T>G SNP (**Table S5D**). We additionally confirmed a subset of the g.27134T>G SNP calls in 16 Coriell samples (n=14 lacking the g.27134T>G SNP, n=2 harboring the g.27134T>G SNP) via *SMN1*-specific PCR and Sanger sequencing (see **Supplementary Methods**), yielding 114 reference samples total (106 unique samples). The NGS results were 100% (114/114) concordant with the reference data (**Figure 4**).

### *FMR1* CGG-repeat analysis

Fragile X syndrome (FXS), the most common cause of inherited intellectual disability, arises from a trinucleotide CGG repeat expansion in the 5’ untranslated region of *FMR1*.^34^ *FMR1* alleles are categorized as normal (NL; 5-44 CGG repeats), intermediate (IM; 45-54 CGG repeats), premutation (PM; 55-200 CGG repeats), and full-mutation (FM; >200 CGG repeats).^35^ We validated our assay using a sample set enriched for expansions of various sizes in both female and male samples. A total of 39 Coriell samples (**Table S5C**) were classified correctly by our assay (**Figure 4C**). Further, the identified CGG repeat allele sizes closely matched the literature consensus sizes (**Figure 4D**).

## DISCUSSION

ECS is gaining widespread clinical adoption—recently receiving support from medical societies^6,36^—because it can provide reliable and affordable risk assessment for many serious recessive and X-linked diseases simultaneously. Genomics technologies like NGS have enabled dramatic growth in ECS panel size without incurring a corresponding rise in testing cost. Coupled with this increase in the achievable panel size, however, is the need to be judicious in panel construction and painstaking in the effort to validate performance. For these reasons, we recently published a systematic process for ECS panel design^14^ and here present both a comprehensive validation study of our updated ECS’s performance and a preliminary analysis of its use on patient samples. Our study of variants in hundreds of genes across hundreds of samples demonstrates high sensitivity, specificity, and accuracy of genotype calls across coding regions and in technically challenging genes (**Figures 3 and 4**).

We introduced panel-wide CNV calling to our NGS-based ECS to maximize the chance of finding couples at risk for children with serious conditions. Though our previous 94-gene ECS could detect six of the most-common CNVs, the updated ECS validated here can identify novel CNVs that span at least one exon in 218 genes. CNVs can be identified in a production workflow via orthogonal technologies (e.g., MLPA) rather than in a single NGS assay (as we do), but MLPA testing is not affordably scalable to hundreds of genes and incurs additional handling steps in the lab that can introduce operator error. Using known-positive samples from biorepositories, retrospectively identified CNV-positive samples, and *in silico* simulations, we demonstrated high sensitivity for novel CNV identification via NGS. Though still preliminary, our analysis of 15,177 samples with the new panel suggests that the ability to identify novel CNVs has a large impact on the efficacy of an ECS, accounting for ∼14% of the total assessed disease risk. We expect that simulation analyses, as we used here, will become increasingly important during NGS-panel validation, where performance needs to be evaluated even when clinical samples are rare or nonexistent.

Not all genes contribute the same amount to the risk assessed by an ECS (**Figure 1**). Indeed, our updated ECS contains more than twice as many genes as the previous version, yet the risk resolved is not twice as great **(Figure 2**). This phenomenon is driven by the disparate incidence of diseases and highlights that it is critical to have high detection rates for the most-common serious conditions, many of which pose screening challenges due to complicated molecular genetics. For several special cases, we have fine-tuned CNV calling to capture single-base differences (SMA), phased and overlapping rearrangements (alpha thalassemia), and very complicated gene conversions (CAH and GBA). In sum, though our risk estimates will become further refined with the addition of more screened patients, we expect the collective risk of these four diseases to rival that of >100 of the least-common diseases on the panel.

Based on the successful validation of the Counsyl Foresight™ Carrier Screen described here, we now broadly perform the test on samples from prospective parents in our clinical laboratory, which is CLIA certified (05D1102604), CAP accredited (7519776), and NYS permitted (8535).

## ACKNOWLEDGEMENTS

We would like to thank our Counsyl colleagues for help and advice: members of the CLIA team, including Thi Tran and Jeanette Wong; Dan Davison; Genevieve Haliburton; Andrew Horn; Eerik Kaseniit; Brandon Lee; Matt Leggett; Jeff Tratner; and Xin Wang.

## REFERENCES

1. Bell CJ, Dinwiddie DL, Miller NA, et al. Carrier testing for severe childhood recessive diseases by next-generation sequencing. Sci Transl Med. 2011;3(65):65ra4.

2. Costa T, Scriver CR, Childs B. The effect of Mendelian disease on human health: a measurement. Am J Med Genet. 1985;21(2):231–242.

3. Kumar P, Radhakrishnan J, Chowdhary MA, Giampietro PF. Prevalence and patterns of presentation of genetic disorders in a pediatric emergency department. Mayo Clin Proc. 2001; 76(8):777–783.

4. Martin JA, Hamilton BE, Osterman MJK, Driscoll AK, Mathews TJ. Births: Final Data for 2015. Natl Vital Stat Rep. 2017;66(1):1.

5. Haque IS, Lazarin GA, Kang HP, Evans EA, Goldberg JD, Wapner RJ. Modeled Fetal Risk of Genetic Diseases Identified by Expanded Carrier Screening. JAMA. 2016;316(7):734–742.

6. Committee Opinion No. 690 Summary: Carrier Screening in the Age of Genomic Medicine. Obstet Gynecol. 2017;129(3):595–596.

7. Azimi M, Schmaus K, Greger V, Neitzel D, Rochelle R, Dinh T. Carrier screening by next-generation sequencing: health benefits and cost effectiveness. Mol Genet Genomic Med. 2016;4(3):292–302.

8. Hallam S, Nelson H, Greger V, et al. Validation for clinical use of, and initial clinical experience with, a novel approach to population-based carrier screening using high-throughput, next-generation DNA sequencing. J Mol Diagn. 2014;16(2):180–189.

9. Umbarger MA, Kennedy CJ, Saunders P, et al. Next-generation carrier screening. Genet Med. 2014;16(2):132–140.

10. Martin J, Asan, Yi Y, et al. Comprehensive carrier genetic test using next-generation deoxyribonucleic acid sequencing in infertile couples wishing to conceive through assisted reproductive technology. Fertil Steril. 2015;104(5):1286–1293.

11. Grody WW, Thompson BH, Gregg AR, et al. ACMG position statement on prenatal/preconception expanded carrier screening. Genet Med. 2013;15(6):482–483.

12. Aradhya S, Lewis R, Bonaga T, et al. Exon-level array CGH in a large clinical cohort demonstrates increased sensitivity of diagnostic testing for Mendelian disorders. Genet Med. 2012;14(6):594–603.

13. Paracchini V, Seia M, Coviello D, et al. Molecular and clinical features associated with CFTR gene rearrangements in Italian population: identification of a new duplication and recurrent deletions. Clin Genet. 2008;73(4):346–352.

14. Beauchamp KA, Muzzey D, Wong KK, et al. Systematic design and comparison of expanded carrier screening panels. Genet Med. June 2017. doi:10.1038/gim.2017.69.

15. Aziz N, Zhao Q, Bry L, et al. College of American Pathologists’ laboratory standards for next-generation sequencing clinical tests. Arch Pathol Lab Med. 2015;139(4):481–493.

16. Rehm HL, Bale SJ, Bayrak-Toydemir P, et al. ACMG clinical laboratory standards for next-generation sequencing. Genet Med. 2013;15(9):733–747.

17. Vysotskaia VS, Hogan GJ, Gould GM, et al. Development and validation of a 36-gene sequencing assay for hereditary cancer risk assessment. PeerJ. 2017;5:e3046.

18. Li H. Aligning sequence reads, clone sequences and assembly contigs with BWA-MEM. arXiv [q-bioGN]. March 2013. http://arxiv.org/abs/1303.3997.

19. Garrison E, Marth G. Haplotype-based variant detection from short-read sequencing. arXiv [q-bioGN]. July 2012. http://arxiv.org/abs/1207.3907.

20. McKenna A, Hanna M, Banks E, et al. The Genome Analysis Toolkit: a MapReduce framework for analyzing next-generation DNA sequencing data. Genome Res. 2010;20(9):1297–1303.

21. Richards S, Aziz N, Bale S, et al. Standards and guidelines for the interpretation of sequence variants: a joint consensus recommendation of the American College of Medical Genetics and Genomics and the Association for Molecular Pathology. Genet Med. 2015;17(5):405–424.

22. Landrum MJ, Lee JM, Riley GR, et al. ClinVar: public archive of relationships among sequence variation and human phenotype. Nucleic Acids Res. 2014;42(Database issue):D980–D985.

23. Kaseniit KE, Theilmann MR, Robertson A, Evans EA, Haque IS. Group Testing Approach for Trinucleotide Repeat Expansion Disorder Screening. Clin Chem. 2016;62(10):1401–1408.

24. 1000 Genomes Project Consortium, Auton A, Brooks LD, et al. A global reference for human genetic variation. Nature. 2015;526(7571):68–74.

25. Chen L, Hadd A, Sah S, et al. An information-rich CGG repeat primed PCR that detects the full range of fragile X expanded alleles and minimizes the need for southern blot analysis. J Mol Diagn. 2010;12(5):589–600.

26. Lim GXY, Yeo M, Koh YY, et al. Validation of a commercially available test that enables the quantification of the numbers of CGG trinucleotide repeat expansion in FMR1 gene. PLoS One. 2017;12(3):e0173279.

27. Clopper CJ, Pearson ES. The Use of Confidence or Fiducial Limits Illustrated in the Case of the Binomial. Biometrika. 1934;26(4):404–413.

28. Zook JM, Chapman B, Wang J, et al. Integrating human sequence data sets provides a resource of benchmark SNP and indel genotype calls. Nat Biotechnol. 2014;32(3):246–251.

29. Lek M, Karczewski KJ, Minikel EV, et al. Analysis of protein-coding genetic variation in 60,706 humans. Nature. 2016;536(7616):285–291.

30. Fokkema IFAC, Taschner PEM, Schaafsma GCP, Celli J, Laros JFJ, den Dunnen JT. LOVD v.2.0: the next generation in gene variant databases. Hum Mutat. 2011;32(5):557–563.

31. Monani UR, Lorson CL, Parsons DW, et al. A single nucleotide difference that alters splicing patterns distinguishes the SMA gene SMN1 from the copy gene SMN2. Hum Mol Genet. 1999;8(7):1177–1183.

32. Nimkarn S, Gangishetti PK, Yau M, New MI. 21-Hydroxylase-Deficient Congenital Adrenal Hyperplasia. University of Washington, Seattle; 2016.

33. Luo M, Liu L, Peter I, et al. An Ashkenazi Jewish SMN1 haplotype specific to duplication alleles improves pan-ethnic carrier screening for spinal muscular atrophy. Genet Med. 2014;16(2):149–156.

34. Saul RA, Tarleton JC. FMR1-Related Disorders. University of Washington, Seattle; 2012.

35. Monaghan KG, Lyon E, Spector EB, American College of Medical Genetics and Genomics. ACMG Standards and Guidelines for fragile X testing: a revision to the disease-specific supplements to the Standards and Guidelines for Clinical Genetics Laboratories of the American College of Medical Genetics and Genomics. Genet Med. 2013;15(7):575–586.

36. Committee Opinion No. 691 Summary: Carrier Screening for Genetic Conditions. Obstet Gynecol. 2017;129(3):597–599.

